# Long-term protein synthesis with PURE in a mesoscale dialysis system

**DOI:** 10.1101/2024.09.09.611992

**Authors:** Laura Roset Julià, Laura Grasemann, Francesco Stellacci, Sebastian J. Maerkl

**Affiliations:** Institute of Materials, School of Engineering, École Polytechnique Fédérale de Lausanne, Lausanne, Switzerland; NCCR Bio-Inspired Materials, École Polytechnique Fédérale de Lausanne, Lausanne, Switzerland; Institute of Bioengineering, School of Engineering, École Polytechnique Fédérale de Lausanne, Lausanne, Switzerland; Global Health Institute, École Polytechnique Fédérale de Lausanne, Lausanne, Switzerland

## Abstract

Cell-free systems are powerful tools in synthetic biology with versatile and wide-ranging applications. However, a significant bottleneck for these systems, particularly the PURE cell-free system, is their limited reaction lifespan and yield. Dialysis offers a promising approach to prolong reaction lifetimes and increase yields, yet most custom dialysis systems require access to sophisticated equipment like 3D printers or microfabrication tools. In this study, we utilized an easy-to-assemble, medium-scale dialysis system for cell-free reactions using commercially available components. By employing dialysis with periodic exchange of the feeding solution, we achieved a protein yield of 1.16 mg/mL GFP in the PURE system and extended protein synthesis for at least 12.5 consecutive days, demonstrating the system’s excellent stability.

## Introduction

Cell-free systems are an ideal chassis for engineering bio-molecular systems due to their versatile, open, and well-defined nature [1, 2, 3, 4, 5]. There are two main types of cell-free systems: lysate-based systems, where the cytoplasm is directly extracted from cells, and the fully recombinant PURE [6] and OnePot PURE system [7, 8]. Current drawbacks of cell-free systems compared to cellular protein expression are a limited reaction time due to a lack of self-regeneration and cellular homeostasis [2], and a limited protein production capacity. Therefore, there is a strong interest in prolonging reaction times to enhance protein yield in cell-free systems. There are generally two approaches to improve protein yield, one is to optimize the composition of the system, and the other is to extend the reaction time by supplying additional energy components and low molecular weight building blocks.

While it is generally the case that lysate systems have higher protein production capacities than PURE, cytoplasmic extracts render lysate systems ill-defined resulting in high batch-to-batch variability [1, 9, 10]. In PURE, all components required for transcription-translation are produced separately, rendering the system well-defined. This is a considerable advantage over lysate systems for a variety of applications including synthetic cell -or therapeutic applications.

To our knowledge, the highest protein yield achieved to date using cell-free systems is 8 mg/mL of protein produced using semi-continuous expression with an optimized lysate system encapsulated in liposomes [11]. Using this lysate formulation, the authors not only improved the yield but also extended protein synthesis to 20 h. The highest protein yield achieved with PURE was reported 10 years ago at 3.8 mg/mL of GFP using a dialysis system. To achieve this yield the authors significantly altered the composition of the PURE system by increasing the concentration of protein and ribosomal components [12].

Other approaches to prolong reaction times in both lysate [13] and PURE systems [14, 15] are based on immobilization or encapsulation strategies. Using these approaches, protein expression was achieved for up to 16 days in PURE [15], and up to 28 days in lysate [13]. These results demonstrated that cell-free systems can sustain protein synthesis for several days in confined and encapsulated systems. However, the hydrogels are labor-intensive to produce and the obtained yield of 200 µg/mL [14] is fairly low. We previously implemented semi-permeable hydrogel membranes in a microfluidic chemostat. Using a commercially available PURE system, we extended protein synthesis at a constant synthesis rate from two to at least 30 h in this microscale dialysis system, and increased overall protein yield by 7-fold [16].

The aforementioned examples require specialized equipment to fabricate microfluidic devices (nL scale) [16, 14, 13, 15] or microscale dialyzer plates (10-50µL) [17, 12]. This limits access to long-lived, high-yield dialysis based cell-free expression systems. For larger scale reactions of around 1 mL, commercially available dialysis devices exist, which often consist of dialysis cups inserted in tubes [18], and thus do not allow monitoring reaction kinetics using standard fluorescent plate readers. Even more problematic are the large reaction volumes, which make these reactions very costly. Commercially available or custom fabricated [19] dialysis plates exist. However, these are either too tall to fit into standard plate reader instruments, or they do not provide physical access for imaging the reaction.

Here we created a simple DIY dialysis system for mesoscale (100 µL) cell-free expression that utilizes commercially available components and can be assembled and used with standard equipment. The main advantages are the open access of the reaction chambers during incubation, allowing feeding solution replenishment, and the possibility for real-time fluorescence monitoring using standard plate readers. Our dialysis system enabled sustained protein synthesis for four days using PURE. By periodically replacing the feeding solution, protein expression was extended to 12.5 days resulting in a protein yield of 1.15 mg/mL. To our knowledge, this represents the longest expression for PURE reported in non-encapsulated systems. Our results highlight the excellent stability of the PURE system and indicate a long protein and ribosome life-time beyond what is currently harnessed in batch reactions and simple dialysis reactions without feeding solution replenishment. We anticipate that this system could be employed in mesoscale protein production in which protein yield is critical. Potential advanced applications in addition to protein production for research purposes could include the de-centralized production of therapeutics [20] as well as the possibility to develop continuous, long-term environmental monitoring [21, 22], or diagnostic systems [23].

## Results

In this work we present a cell-free expression system complemented with a mesoscale dialysis chamber that can be prepared entirely from commercially available components. We used a 24-well plate as the basis for our DIY dialysis system. Each dialysis chamber consists of an independent feeding compartment, and a reaction chamber consisting of the dialysis cup. Dialysis cups have to fulfill two requirements. First, a molecular weight cut-off of 10 kDa is required, as this was shown to be ideal for coupling dialysis to cell-free protein expression [24]. Next, the geometry of the dialysis cup needs to allow the dialysis membrane to be fully immersed in the feeding solution, and the content of the reaction chamber needs to be accessible for imaging. Taking these considerations into account, we chose the Slide-A-Lyzer, 10K, with a 0.1 mL volume as our reaction chamber, which we cut to adjust its size to fit into the well plate (Figure 1A). This setup runs 100 µL cell-free reactions in 1000 µL feeding solution and allows easy exchange of the feeding solution (Figure 1B). We tested this dialysis with lysate reactions (RTS 500 E. Coli HY (Biotechrabbit GmbH)) and PURE reactions (PURExpress (NEB)) combined with a home-made energy solution.

**Figure 1.**
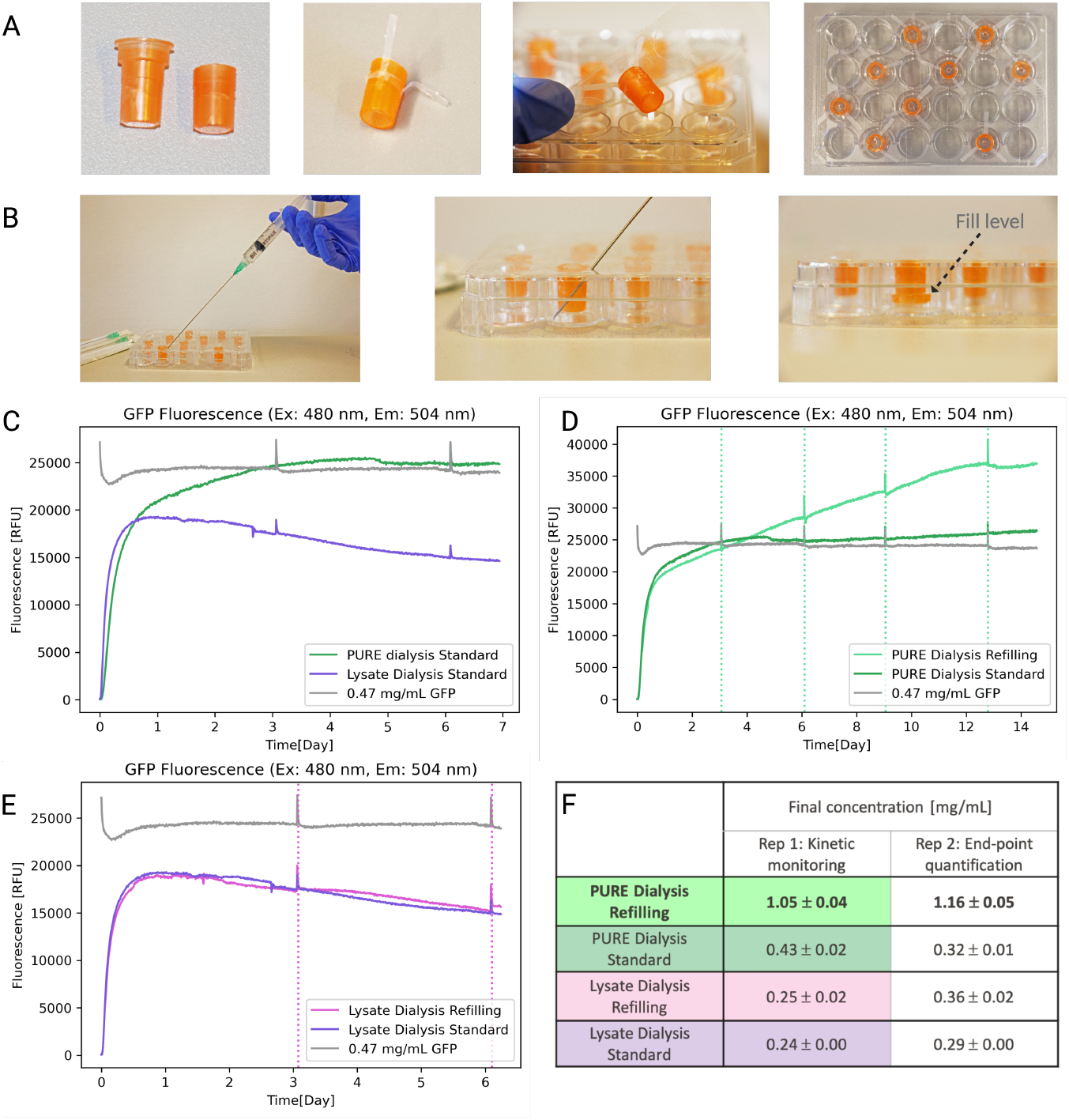
(A) Image of the dialysis cup after and before being cut (left), employed to assemble the dialysis plate (second and third picture), and final appearance of the plate including the plate sealant (right). The cups were distributed randomly across the plate. Theoretically, 24 reactions can be run in parallel. (B) Photographs of the feeding solution refilling procedure. (C) Fluorescence from PURE (dark green) compared to lysate (purple) expressions when the dialysis compartment is not replenished, as well as a reference measurement (grey) of a 0.47 mg/mL solution of GFP in H_2_O, immersed in a feeding compartment containing 1 mL of H_2_O. (D) Fluorescence from PURE reactions with replenishment of the feeding compartment (indicated by vertical dotted lines) in light green, and without replenishment of the feeding compartment, in dark green. (E) Fluorescence from Lysate reactions in dialysis mode, with replenishment of the feeding compartment (indicated by vertical dotted lines) in pink, and without replenishment of the feeding compartment, in purple. (F) Endpoint concentration calculations of the two replicates for each experiment indicating the mean and the standard deviation of technical duplicates for each replicate of the experiment. GFP concentrations were calculated via a GFP calibration curve (Supplementary Figure SI1).

**Figure 1.**
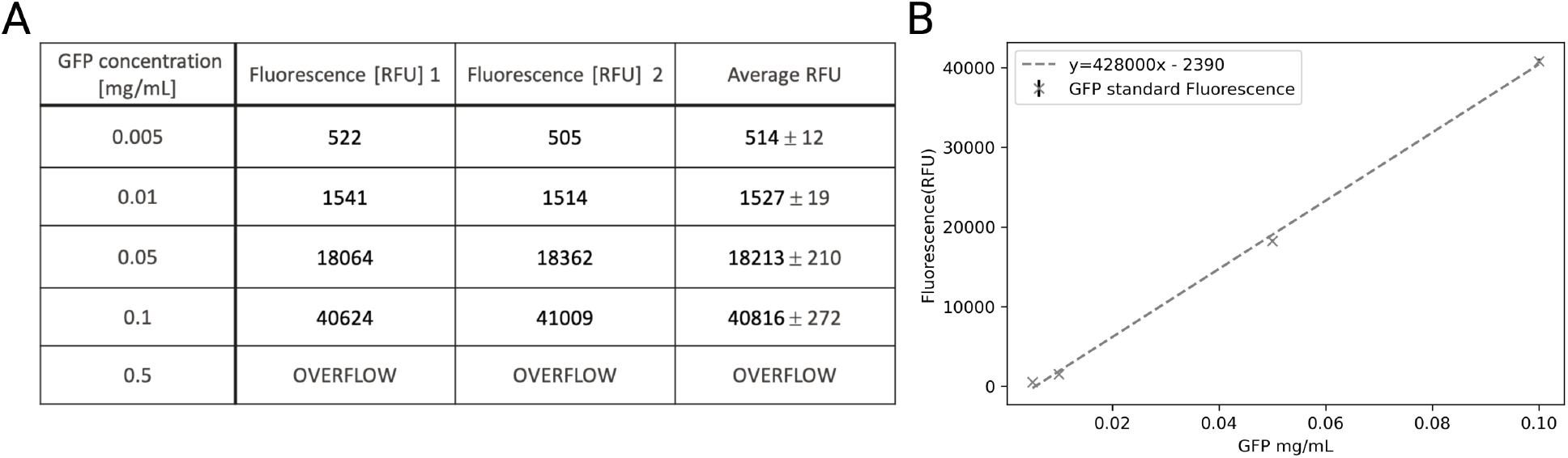
(A) Relative Fluorescence Units (RFU) of the calibration standards, measured in duplicate. The first two columns contain the measured data, while the third column is an average of the two, used to perform the linear regression. (B) Plot of the linear regression (gray dashed line) and the averaged RFU of the calibration standards (marked with a cross), with the measured standard deviation.

**Figure 2.**
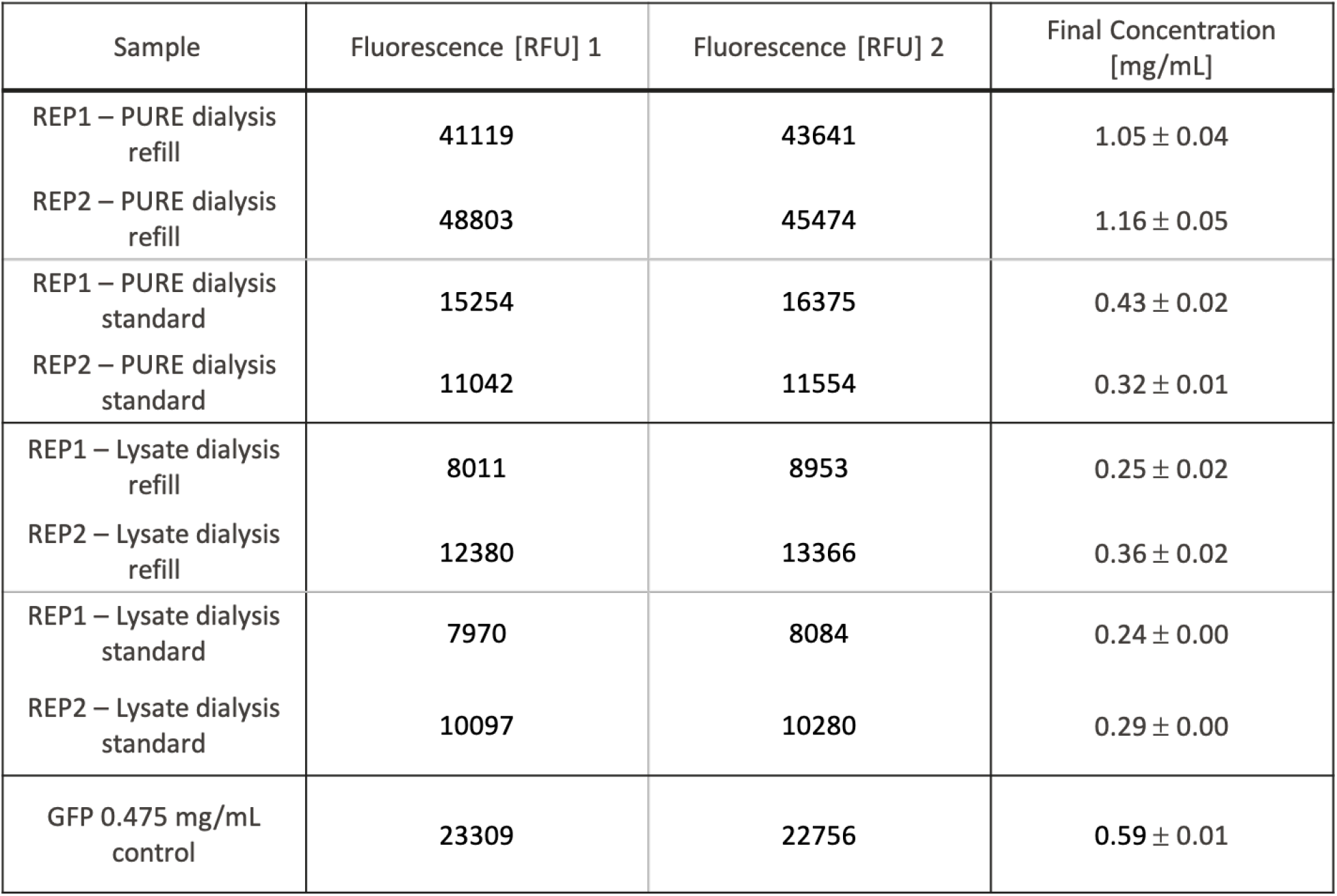
Endpoint measure of the relative fluorescence units (RFU) in technical duplicates (first and second columns) of both replicates for each experiment. The last column is the averaged quantification of the duplicates using the calibration curve presented in Figure 1. The standard deviation results from the technical duplicates.

**Figure 3.**
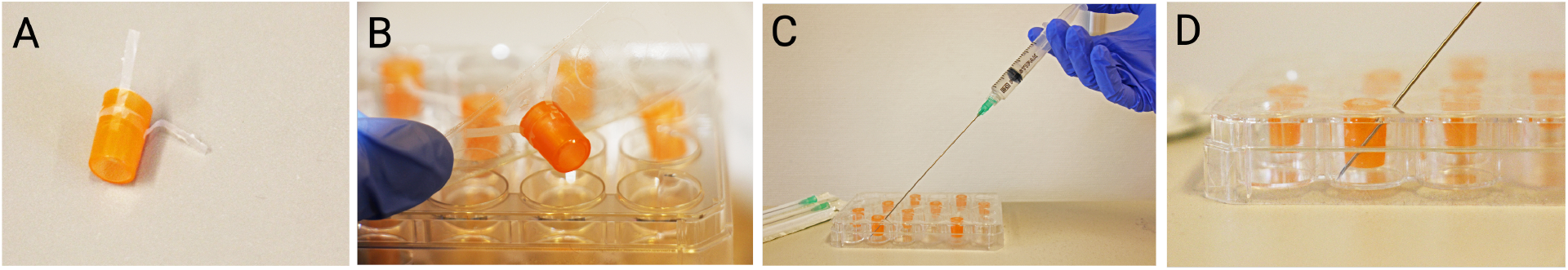
Assembly of the dialysis plate. A and B illustrate the sticking of the cut cups to the plate.

First, we assessed the influence of dialysis on GFP expression in lysate and PURE reactions (Figure 1C). Protein synthesis in a lysate reaction stops after less than one day, while PURE protein synthesis continued for about four days, leading to a higher overall protein yield for the PURE reaction of about 25300 RFU versus 19200 RFU for lysate.

We then set out to investigate, whether replenishing the feeding solution can further prolong cell-free reactions. We replenished the feeding solution every three to four days. Experiments for both PURE and lysate were conducted in the same plate, and each condition was performed in duplicate. Expression of one of the replicates was monitored on the plate reader (Figure 1D,E), while the second replicate was incubated in parallel and used only for end point protein quantification. The PURE reactions were incubated for a total of 14 days and incubation was briefly interrupted four times to exchange the feeding solution for the PURE reactions. Periodic exchange of the feeding solution extended protein synthesis by eight days compared to a reaction without replenishment, leading to active protein synthesis for 12.5 days (Figure 1D). For lysate reactions, the feeding solution was only exchanged during the first two exchanges as protein synthesis did not recover after the first exchange (Figure 1E).

Lastly, total protein expression yield was determined for each sample. Reactions were recovered from the dialysis cups by dilution and re-suspension with a defined volume of water leading to a ten time dilution of the recovered fraction. This ensured re-suspension of potential protein precipitates caused by the long incubation time and the high overall protein concentration. Each solution was then introduced into the plate reader with technical duplicates together with a GFP calibration curve for quantification by fluorescence (Figure 1F and Figures SI1 and SI2). PURE reactions using dialysis without replenishment had an overall yield of 0.43 ± 0.02 mg/mL and 0.32 ± 0.01 mg/mL respectively. Replenishing the feeding solution every 3.5 days resulted in a 2.8-fold increase and a total protein yield of 1.05 ± 0.04 and 1.16 ± 0.05 mg/mL GFP, resulting in mean concentrations of 31.18 mM and 34.49 mM respectively. Protein yields for lysate reactions were considerably lower, and replenishing the feeding solution did not lead to an increase in protein yield. Final protein concentrations in lysate experiments ranged between 0.24 ± 0.00 mg/mL and 0.36 ± 0.02 mg/mL.

## Discussion

In this work, we introduce a simple system for mesoscale cell-free protein expression augmented with dialysis. Using this dialysis system, we extended active protein synthesis in PURExpress from a few hours [6, 7, 8] to around 4 days without exchanging the feeding solution in the feeding compartment. Exchanging the feeding solution every 3-4 days further extended active protein synthesis to 12.5 days, and an overall protein yield of 1 mg/mL. This presents an increase in total protein yield of at least five-fold compared to concentrations of around 150 µg/mL without dialysis [7]. It needs to be mentioned that after 14 days of incubation, precipitate accumulation was observed on the dialysis membranes, although we could not determine at which point precipitate formation started. These precipitates might impede exchange of small molecules across the dialysis membranes, hindering the continued supply of small molecules and the dilution of inhibitory molecules inside the PURE reaction, and could impact fluorescence imaging. It is thus not clear whether cessation of protein synthesis after 12.5 days occurred because of PURE component degradation, or due to obstructed dialysis. Using the same DIY dialysis system did not extend protein synthesis of a lysate-based system and the overall protein yield was substantially lower. We reason that cessation of protein synthesis could be due to degradation of lysate components[10, 18]. Recent findings by Ouyang and coworkers demonstrated active protein synthesis in lysate for 28 days using hydrogel beads [13]. Encapsulation thus seems to prevent those degradation processes and seems to be required for prolonged protein synthesis in lysate systems. Interestingly, PURE sustains a stable synthesis rate without encapsulation for 12.5 days which is comparable to the previously published value of 11-16 days using hydrogel encapsulation [14, 15]. This seems to indicate that the PURE formulation is sufficiently free of proteases, which could negatively impact long-term protein synthesis. Protein expression using our dialysis system resulted in an increase in protein yield of about 5-fold compared to hydrogel based expression [14, 15], rendering the open dialysis system more suitable for applications where high protein concentrations are beneficial, in addition to being simpler to use.

Kazuta and coworkers, have shown that commercial PURE formulations are not optimized for high yield protein expression [12]. It will be interesting to see, what yields can be achieved when combining optimized PURE compositions with simple mesoscale dialysis systems and periodic exchange of solutions. One avenue towards further increasing protein yield may be by reducing protein aggregation, for instance through the addition of chaperones [25].

## Methods

### Preparation of the dialysis plate

The dialysis cups (Slide-A-Lyzer MINI Dialysis Device, 10K MWCO, 0.1 mL) were cut below the rim to fit into the 24 well plate (Nunc, Thermo Fisher). The cups were then adhered to a transparent plate sealant (SealPlate film Z369659) while still empty, by securing them with three thin strips of tape. An additional strip of tape was wrapped around the entire perimeter of the cup for further reinforcement, and the cup-containing sealant was arranged on top of the plate ensuring the dialysis cup was centered in the well (Figure 1A). 1 mL of feeding solution (see below) was introduced into the feeding compartment by punching the seal with a needle. Subsequently, 100 *µ*L of the reaction solution (see below) were introduced into the dialysis cup. After the assembly of all reactions, plasmid DNA (see below) was introduced into each dialysis cup to initiate the reactions. The plate was then sealed with an additional layer of plate sealant and placed in the plate reader. The plate was incubated at 32°C, monitoring the fluorescence over time (Excitation 480 nm, Emission: 504 nm).

To replenish the feeding solution, the plate was removed from the plate reader. Subsequently, the seal was perforated with a syringe and the spent feeding solution was aspirated. Using a fresh syringe, the feeding compartment was then replenished with fresh solution, the plate was sealed with another layer of transparent plate sealant, and the plate was inserted back into the plate reader for another round of incubation (Figure 1B). A total volume of 1000 µL of feeding solution in the feeding compartment was sufficient to entirely immerse the dialysis membrane in solution.

### Energy and feeding solution preparation for PURE

The energy and feeding solution was prepared as previously published [8, 16] at 2.5x, but omitting tRNAs. The 2.5x energy and feeding solutions contained 125 mM HEPES, 250 mM potassium glutamate, 29.5 mM magnesium acetate, 5 mM ATP and GTP respectively, 2.5 mM UTP and CTP respectively, 50 mM creatine phosphate, 2.5 mM TCEP, 0.05 mM folinic acid, 5 mM spermidine, and 0.75 mM of each amino acid. The solution was used at 2.5x concentration in the feeding solution, and was added as a 1x energy solution in the PURE reaction in the reaction compartment, supplemented with tRNAs.

The tRNAs were purified from *E. coli* BL21, slightly adapted from a previously described protocol [26]. Briefly, a culture of *E. coli* BL21 cells was grown at 37°C for 6 h, and cells were harvested by centrifugation. The cell pellet was weighed and resuspended in five times the weight of the pellet in resuspension buffer (10 mM HEPES, 10 mM MgCl_2_, pH 7.2) e.g. 5 g cell pellet was resuspended in 25 mL resuspention buffer. The same amount of equilibrated phenol (Invitrogen) was added to achieve a 1:1 v/v ratio. The solution was mixed and incubated for 30 min at 4°C on a lab rotisserie. Phases were separated by centrifugation for 10 min at 4000 x g and the aqueous phase was transferred to a new tube. Ultrapure isopropanol was added to achieve a 1:1 v/v ratio and the solution was incubated at -20°C over night. Nucleic acids were precipitated by centrifugation for 10 min at 4°C and 4000 x g. The pellet was resuspended in lithium buffer (0.8 M LiCl, 0.8 M NaCl), and repelleted with another centrifugation step using the previous conditions. The supernatant was subsequently transferred to a new tube, isopropanol was added to achieve a v/v ratio of 1:1, followed by a precipitation step using the previous centrifugation parameters. The pellet was dissolved in ultrapure ethanol and subsequently pelleted for three rounds. After the third round, the pellet was dried with nitrogen and subsequently dissolved in as little final buffer volume as possible (40 mM NaCl, 10 mM HEPES in ultrapure water), typically resulting in a final concentration of around 25 mg/mL. 1 µL of protector RNase inhibitor was added to approx 200 µL of tRNA solution, and tRNAs were stored at -80°C until further use.

### Experimental setup for PURE expression with dialysis

For PURE protein expression, 1x PURExpress (NEB) solution B was supplemented with RNase inhibitor (2 U/µL), mScarlet, 10 mM TCEP, 0.47 mg/mL ampicillin, 1.5 mg/mL tRNAs, and 1x energy solution. The energy solution A from the commercially available kit was replaced by home-made energy solution, as the beta-mercaptoethanol present in solution A is less stable than the TCEP used in the home made solution, and was previously suspected to limit the life time of PURE reactions [16]. The solution was assembled as a master mix for all reactions at a total reaction volume of 420 µL. 100 µL PURE solution were distributed to each dialysis cup, and the reaction was initiated by adding 5 nM of plasmid DNA. The feeding solution at 2.5x was supplemented with 2.5 mg/mL of ampicillin and applied to the feeding compartment. A 2.5 x concentration has previously shown to be optimal for dialysis reactions in PURE [16]. It needs to be noted that, unlike in previous publications, the tRNAs were added solely to the reaction compartment, as their molecular weight would prevent them from diffusing via the 10 kDa cut-off dialysis membrane. Plasmid template DNA was used at a concentration of 5 nM.

### Experimental setup for Lysate expression with dialysis

Lysate reactions were assembled following the supplier’s protocol (BR1400201 RTS 500 Proteo-Master E. coli HY Kit -Biotechrabbit Gmbh), omitting the dialysis reaction device. The synthesis reaction was assembled as a master mix for all reactions in a total volume of 1030.5 µL using only freshly reconstituted components. The master mix included 525.5 µL of Lysate, 225 µL of Reaction Mix, 250 µL of Amino Acids and 30 µL of Methionine. The solution was mixed, and 100 µL were added into each dialysis cup. Remaining solution was either stored or directly used for further experiments. Plasmid DNA was added at a final concentration of 3.4 nM, as recommended by the supplier. The Feeding Mix was assembled by adding 2.65 mL of Amino Acids and 300 µL of Methionine.

### 0.1 DNA preparation

For all experiments, plasmid DNA encoding muGFP was used. The plasmid is a pET 29b(+) plasmid, and was purified from *E. coli* 10-beta (NEB) cells using a Zymo Miniprep kit according to the supplier’s instructions. DNA elution was performed in water instead of elution buffer, as the latter contains EDTA, which complexes Mg ions and is thus detrimental for cell-free reactions.

### Experimental setup for the GFP control

GFP in water was used for the control well during the experiment, and for the mass calibration curve used for the endpoint quantification of the samples. It was obtained as follows. The same plasmid (section 0.1) used for the experiments was transformed into BL21(DE3) (Lucigen) cells and plated. A colony of cells was grown overnight in LB media supplemented in kanamycin. For every liter of production culture, 20 mL of overnight culture were inoculated into Autoinduction TB (Formedium) media and grown at 37°C. A total of 12 L culture was grown in batches of 2 L in 5 L flasks. After 4-5 hours or an OD600 of *>*0.8, the incubator temperature was changed to 18°C. The cultures were incubated overnight for at least 18 hours. Cell pellets were harvested and stored at -20°C until further use. Cell pellets were resuspended in Buffer A (700 mM NaCl, 20 mM HEPES 7.5), supplemented with glycerol to 10% v/v and 10 µL of Turbonuclease, then lysed by sonication. Sonication was performed for 2 minutes and 30 seconds in total, in pulses of 10 seconds on and 10 seconds off. For every 2 L of pellet, the approximate volume of lysate was 50 mL. Lysates were clarified by centrifugation, filtered through a 0.45 um filter and supplemented with 25 mM imidazole. The sample was loaded onto a 25 mL NiNTA column (Cytiva or ProteinArk) and the protein was eluted on a gradient of 5-100% Buffer B (700 mM NaCl, 500 mM imidazole, 20 mM HEPES 7.5) on an AKTA system. Pooled eluted fractions were dialyzed twice in 5 L of MilliQ water. The final concentration of the GFP solution was 1.87 mg/mL determined using a nanodrop at 280 nm.

The obtained solution was diluted to 0.47 mg/mL and introduced in the dialysis cup. The feeding compartment was filled with 1 mL of water. During the exchanges of the experiment reaction, the dialysis chamber was replenished with fresh water as well. The fluorescence was the monitored throughout the incubation period (Figure 1).

### Endpoint measurements

After 14 days of incubation, the plate was removed from the plate reader. The solutions in the reaction compartment were recovered by aspirating the content with a pipette, followed by several washing steps with water to recover all protein. This resulted in a final volume of 1 mL for each reaction, representing a 1:10 dilution. Duplicates of 40 µL for each reaction chamber were introduced in a 384 well plate for fluorescence measurement on the plate reader.

For the GFP calibration curve, the muGFP solution (see above) was diluted to 0.5, 0.1, 0.05, 0.01, and 0.005 mg/mL and applied to the same 384 plate in duplicates. The plate was centrifuged for 30 seconds at 10 000 x g and introduced into the plate reader. Readings were performed after 10 seconds shaking, and at room temperature. The calibration curve was obtained from a linear regression of the fluorescence counts averaged for the duplicates (Supplementary Figure SI1). Sample concentrations were calculated from the measured RFU and the slope and intercept of the calibration curve. The obtained concentrations were averaged within the technical duplicates and the standard deviation was calculated (Supplementary Figure SI2).

The GFP control of our DIY dialysis system was used to assess the accuracy of the endpoint quantification method. By performing the same recovery method of the control well, we included duplicates of the GFP control to obtain the endpoint fluorescence of the latter, which allowed us to back-calculate the concentration. We obtained a value of 0.594 ± 0.009 mg/mL, indicating that the quantification method has an accuracy of about 25%.

## Author Contributions

L.R. and L.G. contributed equally and performed experiments. L.R., L.G., F.S. and S.J.M. designed experiments, analyzed data, and wrote the manuscript.

## Acknowledgements

The authors would like to thank Matt Cummins for sharing his protocol for tRNA preparation, and Kelvin Lau for helpful input and comments as well as purifying the GFP used for the controls. Thanks to the lab members from LBNC and SuNMIL for helpful input and comments.

## Supplementary Information

### plasmid sequence

CCGGTGATGCCGGCCACGATGCGTCCGGCGTAGAGGATCGAGATCGATCTCGATCCCGCGAAATTAATACGACTCACTATAGGGGAATTGTGAGCGGATAACAATTCC CCTCTAGAAATAATTTTGTTTAACTTTAAGAAGGAGATATACATATGAAACATCATCATCACCACCATCATCACCACCACCATCACCATCATGGCGCCTCTATGGATTA CAAAGACCATGATGGTGATTACAAGGATCATGATATTGATTATAAAGATGATGATGATAAAGGTTCCGGGTCTGGTGAAAACCTCTACTTTCAAGGGTCGGGATCCAT GGTTTCCAAAGGAGAAGAACTGTTTACCGGTGTTGTACCAATTCTCGTAGAACTCGATGGAGATGTAAACGGGCATAAATTTTCAGTGCGCGGCGAGGGCGAAGGAGA TGCCACAAACGGCAAACTGACCCTTAAATTTATTTGCACGACCGGCAAATTACCAGTTCCTTGGCCTACGCTGGTCACCACGCTCACCTATGGGGTATTATGCTTTAGCC GCTATCCGGATCACATGAAACGCCATGATTTCTTTAAAAGTGCTATGCCAGAAGGTTATGTACAGGAACGCACGATTAGCTTTAAAGATGATGGGACGTATAAAACCC GCGCCGAGGTAAAATTTGAAGGAGATACCTTAGTAAACCGCATTGAACTCAAGGGGATTGATTTTAAAGAGGACGGTAACATTCTGGGTCATAAACTTGAGTACAACT TTAACTCACACAACGTTTACATTACCGCGGATAAACAGAAGAACGGTATTAAAGCGTACTTTAAGATTCGCCATAACGTCGAAGATGGCAGTGTTCAGCTGGCCGATC ATTATCAGCAGAACACGCCGATTGGCGATGGCCCTGTTTTGTTACCGGATAACCATTATTTATCGACTCAGAGCGTCTTAAGTAAAGATCCAAACGAGAAACGCGATC ACATGGTTCTCTTAGAAGATGTTACCGCCGCCGGCATTACACATGGCATGGATGAACTGTATAAATGATAGGCGGCCGCAGGACTGAATGATATTTTCGAAGCACAAA AGATTGAGTGGCATGAAGCTAGCGAGAATTTGTATTTTCAAGGTAGTGCTTGGTCGCACCCTCAATTCGAAAAGGGCGGCGGTAGTGGCGGTGGTTCAGGCGGTTCCGC GTGGAGTCACCCGCAATTCGAGAAAGGCGCTTGATAGCTCGAGCACCACCACCACCACCACTGAGATCCGGCTGCTAACAAAGCCCGAAAGGAAGCTGAGTTGGCTG CTGCCACCGCTGAGCAATAACTAGCATAACCCCTTGGGGCCTCTAAACGGGTCTTGAGGGGTTTTTTGCTGAAAGGAGGAACTATATCCGGATTGGCGAATGGGACGC GCCCTGTAGCGGCGCATTAAGCGCGGCGGGTGTGGTGGTTACGCGCAGCGTGACCGCTACACTTGCCAGCGCCCTAGCGCCCGCTCCTTTCGCTTTCTTCCCTTCCTTTC TCGCCACGTTCGCCGGCTTTCCCCGTCAAGCTCTAAATCGGGGGCTCCCTTTAGGGTTCCGATTTAGTGCTTTACGGCACCTCGACCCCAAAAAACTTGATTAGGGTGAT GGTTCACGTAGTGGGCCATCGCCCTGATAGACGGTTTTTCGCCCTTTGACGTTGGAGTCCACGTTCTTTAATAGTGGACTCTTGTTCCAAACTGGAACAACACTCAACCC TATCTCGGTCTATTCTTTTGATTTATAAGGGATTTTGCCGATTTCGGCCTATTGGTTAAAAAATGAGCTGATTTAACAAAAATTTAACGCGAATTTTAACAAAATATTAAC GCTTACAATTTAGGTGGCACTTTTCGGGGAAATGTGCGCGGAACCCCTATTTGTTTATTTTTCTAAATACATTCAAATATGTATCCGCTCATGAATTAATTCTTAGAAAAA CTCATCGAGCATCAAATGAAACTGCAATTTATTCATATCAGGATTATCAATACCATATTTTTGAAAAAGCCGTTTCTGTAATGAAGGAGAAAACTCACCGAGGCAGTTC CATAGGATGGCAAGATCCTGGTATCGGTCTGCGATTCCGACTCGTCCAACATCAATACAACCTATTAATTTCCCCTCGTCAAAAATAAGGTTATCAAGTGAGAAATCAC CATGAGTGACGACTGAATCCGGTGAGAATGGCAAAAGTTTATGCATTTCTTTCCAGACTTGTTCAACAGGCCAGCCATTACGCTCGTCATCAAAATCACTCGCATCAAC CAAACCGTTATTCATTCGTGATTGCGCCTGAGCGAGACGAAATACGCGATCGCTGTTAAAAGGACAATTACAAACAGGAATCGAATGCAACCGGCGCAGGAACACTG CCAGCGCATCAACAATATTTTCACCTGAATCAGGATATTCTTCTAATACCTGGAATGCTGTTTTCCCGGGGATCGCAGTGGTGAGTAACCATGCATCATCAGGAGTACG GATAAAATGCTTGATGGTCGGAAGAGGCATAAATTCCGTCAGCCAGTTTAGTCTGACCATCTCATCTGTAACATCATTGGCAACGCTACCTTTGCCATGTTTCAGAAAC AACTCTGGCGCATCGGGCTTCCCATACAATCGATAGATTGTCGCACCTGATTGCCCGACATTATCGCGAGCCCATTTATACCCATATAAATCAGCATCCATGTTGGAATT TAATCGCGGCCTAGAGCAAGACGTTTCCCGTTGAATATGGCTCATAACACCCCTTGTATTACTGTTTATGTAAGCAGACAGTTTTATTGTTCATGACCAAAATCCCTTAA CGTGAGTTTTCGTTCCACTGAGCGTCAGACCCCGTAGAAAAGATCAAAGGATCTTCTTGAGATCCTTTTTTTCTGCGCGTAATCTGCTGCTTGCAAACAAAAAAACCACC GCTACCAGCGGTGGTTTGTTTGCCGGATCAAGAGCTACCAACTCTTTTTCCGAAGGTAACTGGCTTCAGCAGAGCGCAGATACCAAATACTGTCCTTCTAGTGTAGCCG TAGTTAGGCCACCACTTCAAGAACTCTGTAGCACCGCCTACATACCTCGCTCTGCTAATCCTGTTACCAGTGGCTGCTGCCAGTGGCGATAAGTCGTGTCTTACCGGGTT GGACTCAAGACGATAGTTACCGGATAAGGCGCAGCGGTCGGGCTGAACGGGGGGTTCGTGCACACAGCCCAGCTTGGAGCGAACGACCTACACCGAACTGAGATACC TACAGCGTGAGCTATGAGAAAGCGCCACGCTTCCCGAAGGGAGAAAGGCGGACAGGTATCCGGTAAGCGGCAGGGTCGGAACAGGAGAGCGCACGAGGGAGCTTCC AGGGGGAAACGCCTGGTATCTTTATAGTCCTGTCGGGTTTCGCCACCTCTGACTTGAGCGTCGATTTTTGTGATGCTCGTCAGGGGGGCGGAGCCTATGGAAAAACGCC AGCAACGCGGCCTTTTTACGGTTCCTGGCCTTTTGCTGGCCTTTTGCTCACATGTTCTTTCCTGCGTTATCCCCTGATTCTGTGGATAACCGTATTACCGCCTTTGAGTGAG CTGATACCGCTCGCCGCAGCCGAACGACCGAGCGCAGCGAGTCAGTGAGCGAGGAAGCGGAAGAGCGCCTGATGCGGTATTTTCTCCTTACGCATCTGTGCGGTATTT CACACCGCAATGGTGCACTCTCAGTACAATCTGCTCTGATGCCGCATAGTTAAGCCAGTATACACTCCGCTATCGCTACGTGACTGGGTCATGGCTGCGCCCCGACACC CGCCAACACCCGCTGACGCGCCCTGACGGGCTTGTCTGCTCCCGGCATCCGCTTACAGACAAGCTGTGACCGTCTCCGGGAGCTGCATGTGTCAGAGGTTTTCACCGTC ATCACCGAAACGCGCGAGGCAGCTGCGGTAAAGCTCATCAGCGTGGTCGTGAAGCGATTCACAGATGTCTGCCTGTTCATCCGCGTCCAGCTCGTTGAGTTTCTCCAGA AGCGTTAATGTCTGGCTTCTGATAAAGCGGGCCATGTTAAGGGCGGTTTTTTCCTGTTTGGTCACTGATGCCTCCGTGTAAGGGGGATTTCTGTTCATGGGGGTAATGAT ACCGATGAAACGAGAGAGGATGCTCACGATACGGGTTACTGATGATGAACATGCCCGGTTACTGGAACGTTGTGAGGGTAAACAACTGGCGGTATGGATGCGGCGGG ACCAGAGAAAAATCACTCAGGGTCAATGCCAGCGCTTCGTTAATACAGATGTAGGTGTTCCACAGGGTAGCCAGCAGCATCCTGCGATGCAGATCCGGAACATAATG GTGCAGGGCGCTGACTTCCGCGTTTCCAGACTTTACGAAACACGGAAACCGAAGACCATTCATGTTGTTGCTCAGGTCGCAGACGTTTTGCAGCAGCAGTCGCTTCACG TTCGCTCGCGTATCGGTGATTCATTCTGCTAACCAGTAAGGCAACCCCGCCAGCCTAGCCGGGTCCTCAACGACAGGAGCACGATCATGCGCACCCGTGGGGCCGCCA TGCCGGCGATAATGGCCTGCTTCTCGCCGAAACGTTTGGTGGCGGGACCAGTGACGAAGGCTTGAGCGAGGGCGTGCAAGATTCCGAATACCGCAAGCGACAGGCCG ATCATCGTCGCGCTCCAGCGAAAGCGGTCCTCGCCGAAAATGACCCAGAGCGCTGCCGGCACCTGTCCTACGAGTTGCATGATAAAGAAGACAGTCATAAGTGCGGC GACGATAGTCATGCCCCGCGCCCACCGGAAGGAGCTGACTGGGTTGAAGGCTCTCAAGGGCATCGGTCGAGATCCCGGTGCCTAATGAGTGAGCTAACTTACATTAAT TGCGTTGCGCTCACTGCCCGCTTTCCAGTCGGGAAACCTGTCGTGCCAGCTGCATTAATGAATCGGCCAACGCGCGGGGAGAGGCGGTTTGCGTATTGGGCGCCAGGGT GGTTTTTCTTTTCACCAGTGAGACGGGCAACAGCTGATTGCCCTTCACCGCCTGGCCCTGAGAGAGTTGCAGCAAGCGGTCCACGCTGGTTTGCCCCAGCAGGCGAAAA TCCTGTTTGATGGTGGTTAACGGCGGGATATAACATGAGCTGTCTTCGGTATCGTCGTATCCCACTACCGAGATGTCCGCACCAACGCGCAGCCCGGACTCGGTAATGG CGCGCATTGCGCCCAGCGCCATCTGATCGTTGGCAACCAGCATCGCAGTGGGAACGATGCCCTCATTCAGCATTTGCATGGTTTGTTGAAAACCGGACATGGCACTCCA GTCGCCTTCCCGTTCCGCTATCGGCTGAATTTGATTGCGAGTGAGATATTTATGCCAGCCAGCCAGACGCAGACGCGCCGAGACAGAACTTAATGGGCCCGCTAACAG CGCGATTTGCTGGTGACCCAATGCGACCAGATGCTCCACGCCCAGTCGCGTACCGTCTTCATGGGAGAAAATAATACTGTTGATGGGTGTCTGGTCAGAGACATCAAG AAATAACGCCGGAACATTAGTGCAGGCAGCTTCCACAGCAATGGCATCCTGGTCATCCAGCGGATAGTTAATGATCAGCCCACTGACGCGTTGCGCGAGAAGATTGTG CACCGCCGCTTTACAGGCTTCGACGCCGCTTCGTTCTACCATCGACACCACCACGCTGGCACCCAGTTGATCGGCGCGAGATTTAATCGCCGCGACAATTTGCGACGGC GCGTGCAGGGCCAGACTGGAGGTGGCAACGCCAATCAGCAACGACTGTTTGCCCGCCAGTTGTTGTGCCACGCGGTTGGGAATGTAATTCAGCTCCGCCATCGCCGCTT CCACTTTTTCCCGCGTTTTCGCAGAAACGTGGCTGGCCTGGTTCACCACGCGGGAAACGGTCTGATAAGAGACACCGGCATACTCTGCGACATCGTATAACGTTACTGG TTTCACATTCACCACCCTGAATTGACTCTCTTCCGGGCGCTATCATGCCATACCGCGAAAGGTTTTGCGCCATTCGATGGTGTCCGGGATCTCGACGCTCTCCCTTATGCG ACTCCTGCATTAGGAAGCAGCCCAGTAGTAGGTTGAGGCCGTTGAGCACCGCCGCCGCAAGGAATGGTGCATGCAAGGAGATGGCGCCCAACAGTCCCCCGGCCACG GGGCCTGCCACCATACCCACGCCGAAACAAGCGCTCATGAGCCCGAAGTGGCGAGCCCGATCTTCCCCATCGGTGATGTCGGCGATATAGGCGCCAGCAACCGCACC TGTGGCG

## References

[1] Nadanai Laohakunakorn, Laura Grasemann, Barbora Lavickova, Grégoire Michielin, Amir Shahein, Zoe Swank, and Sebastian Josef Maerkl. Bottom-up construction of complex biomolecular systems with cell-free synthetic biology. Frontiers in Bioengineering and Biotechnology, 8:213, mar 2020.

[2] Nadanai Laohakunakorn. Cell-Free Systems: A Proving Ground for Rational Biodesign. Frontiers in Bioengineering and Biotechnology, 8:788, jul 2020.

[3] David Garenne, Matthew C. Haines, Eugenia F. Romantseva, Paul Freemont, Elizabeth A. Strychalski, and Vincent Noireaux. Cell-free gene expression. Nature Reviews Methods, 1(1):49, 2021.

[4] Jessica G. Perez, Jessica C. Stark, and Michael C. Jewett. Cell-free synthetic biology: Engineering beyond the cell. Cold Spring Harbor Perspectives in Biology, 8(12):a023853, ec 2016.

[5] Vincent Noireaux and Allen P. Liu. The New Age of Cell-Free Biology. 22(Volume 22, 2020):51–77, jun 2020.

[6] Yoshihiro Shimizu, Akio Inoue, Yukihide Tomari, Tsutomu Suzuki, Takashi Yokogawa, Kazuya Nishikawa, and Takuya Ueda. Cell-free translation reconstituted with purified components. Nature Biotechnology, 19(8):751–755, 2001.

[7] Barbora Lavickova and Sebastian J. Maerkl. A Simple, Robust, and Low-Cost Method to Produce the PURE Cell-Free System. ACS Synthetic Biology, 2019.

[8] Laura Grasemann, Barbora Lavickova, M. Carolina Elizondo-Cantú, and Sebastian J. Maerkl. Onepot pure cell-free system. Journal of Visualized Experiments, 2021(172):62625, jun 2021.

[9] David Garenne, Chase L. Beisel, and Vincent Noireaux. Characterization of the all-e. coli transcription-translation system mytxtl by mass spectrometry. Rapid Communications in Mass Spectrometry, 33(11):1036–1048, 2019.

[10] Daniel Foshag, Erik Henrich, Ekkehard Hiller, Miriam Schäfer, Christian Kerger, Anke Burger-Kentischer, Irene Diaz-Moreno, Sofía M. García-Mauriño, Volker Dötsch, Steffen Rupp, and Frank Bernhard. The E. coli S30 lysate proteome: A prototype for cell-free protein production. New Biotechnology, 40:245–260, 2018.

[11] D Garenne, S Thompson, A Brisson Synthetic …, and Undefined 2021. The all-E. coliTXTL toolbox 3.0: new capabilities of a cell-free synthetic biology platform. Synthetic Biology, 6(1):ysab017, 2021.

[12] Yasuaki Kazuta, Tomoaki Matsuura, Norikazu Ichihashi, and Tetsuya Yomo. Synthesis of milligram quantities of proteins using a reconstituted in vitro protein synthesis system. Journal of Bioscience and Bioengineering, 118(5):554–557, nov 2014.

[13] Xiaofei Ouyang, Xiaoyu Zhou, Sze Nga Lai, Qi Liu, and Bo Zheng. Immobilization of Proteins of Cell Extract to Hydrogel Networks Enhances the Longevity of Cell-Free Protein Synthesis and Supports Gene Networks. ACS Synthetic Biology, 10(4):749–755, apr 2021.

[14] X Zhou, H Wu, M Cui, SN Lai, B Zheng Chemical Science, and Undefined 2018. Long-lived protein expression in hydrogel particles: towards artificial cells. Chemical Science, 9(18):4275–4279, 2018.

[15] Sze Nga Lai, Xiaoyu Zhou, Xiaofei Ouyang, Hui Zhou, Yujie Liang, Jiang Xia, and Bo Zheng. Artificial Cells Capable of Long-Lived Protein Synthesis by Using Aptamer Grafted Polymer Hydrogel. ACS Synthetic Biology, 9(1):76–83, jan 2020.

[16] Barbora Lavickova, Laura Grasemann, and Sebastian J. Maerkl. Improved Cell-Free Transcription-Translation Reactions in Microfluidic Chemostats Augmented with Hydrogel Membranes for Continuous Small Molecule Dialysis. ACS Synthetic Biology, 11(12):4134–4141, ec 2022.

[17] K Jackson, T Kanamori, and T Ueda. Protein synthesis yield increased 72 times in the cell-free PURE system. Integrative Biology, 6(8):781–788, 2014.

[18] Daniel Schwarz, Friederike Junge, Florian Durst, Nadine Frölich, Birgit Schneider, Sina Reckel, Solmaz Sobhanifar, Volker Dötsch, and Frank Bernhard. Preparative scale expression of membrane proteins in Escherichia coli-based continuous exchange cell-free systems. Nature Protocols, 2(11):2945–2957, 2007.

[19] Masaaki Aoki, Takayoshi Matsuda, Yasuko Tomo, Yukako Miyata, Makoto Inoue, Takanori Kigawa, and Shigeyuki Yokoyama. Automated system for high-throughput protein production using the dialysis cell-free method. Protein Expression and Purification, 68(2):128–136, 2009.

[20] Keith Pardee, Shimyn Slomovic, Peter Q. Nguyen, Jeong Wook Lee, Nina Donghia, Devin Burrill, Tom Ferrante, Fern R. McSorley, Yoshikazu Furuta, Andyna Vernet, Michael Lewandowski, Christopher N. Boddy, Neel S. Joshi, and James J. Collins. Portable, On-Demand Biomolecular Manufacturing. Cell, 167(1):248–259.e12, sep 2016.

[21] Walter Thavarajah, Adam D. Silverman, Matthew S. Verosloff, Nancy Kelley-Loughnane, Michael C. Jewett, and Julius B. Lucks. Point-of-use detection of environmental fluoride via a cell-free riboswitch-based biosensor. ACS Synthetic Biology, 9(1):10–18, 2020. PMID: 31829623.

[22] Adam D. Silverman, Umut Akova, Khalid K. Alam, Michael C. Jewett, and Julius B. Lucks. Design and optimization of a cell-free atrazine biosensor. ACS Synthetic Biology, 9(3):671–677, 2020. PMID: 32078765.

[23] Aidan Tinafar, Katariina Jaenes, and Keith Pardee. Synthetic Biology Goes Cell-Free. BMC Biology 2019 17:1, 17(1):1–14, aug 2019.

[24] Hoang Thanh Nguyen, Morgan Massino, Camille Keita, and Jean Baptiste Salmon. Microfluidic dialysis using photo-patterned hydrogel membranes in PDMS chips. Lab on a Chip, 20(13):2383–2393, jul 2020.

[25] Tatsuya Niwa, Takashi Kanamori, Takuya Ueda, and Hideki Taguchi. Global analysis of chaperone effects using a reconstituted cell-free translation system. Proceedings of the National Academy of Sciences of the United States of America, 109(23):8937–8942, jun 2012.

[26] E Cayama, A Yépez, F Rotondo, E Bandeira, AC Ferreras, and FJ Triana-Alonso. New chromatographic and biochemical strategies for quick preparative isolation of tRNA. Nucleic Acids Research, 28(12):e64–e64, 2000.

